# Cell volume homeostatically controls the rDNA repeat copy number and rRNA synthesis rate in yeast

**DOI:** 10.1101/841304

**Authors:** José E. Pérez-Ortín, Adriana Mena, Marina Barba-Aliaga, Rebeca Alonso-Monge, Abhyudai Singh, Sebastián Chávez, José García-Martínez

## Abstract

The adjustment of transcription and translation rates to variable needs is of utmost importance for the fitness and survival of living cells. We have previously shown that the global transcription rate for RNA polymerase II is regulated differently in cells presenting symmetrical or asymmetrical cell division. The budding yeast *Saccharomyces cerevisiae* adopts a particular strategy to avoid that the smaller daughter cells increase their total mRNA concentration with every generation. The global mRNA synthesis rate lowers with a growing cell volume, but global mRNA stability increases. In this paper, we address what the solution is to the same theoretical problem for the RNA polymerase I synthesis rate. We find that the RNA polymerase I synthesis rate strictly depends on the copy number of its 35S rRNA gene. For cells with larger cell sizes, such as a mutant *cln3* strain, the rDNA repeat copy number is increased by a mechanism based on a feed-back mechanism in which Sir2 histone deacetylase homeostatically controls the amplification of the rRNA genes at the rDNA locus in a volume-dependent manner.

## Introduction

Eukaryotic cells have distributed the transcription work in its nuclei among three different RNA polymerases (RNA pol). RNA pol II transcribes most genes, including those that encode proteins. RNA pol III transcribes about several hundreds of genes (mostly tRNAs and 5S rRNA genes). RNA pol I, specialised in transcribing only one gene, the large rRNA 35-47S (depending on the particular eukaryote) gene is, however, the biggest consumer of ribonucleotides given the need for a vast amount of this precursor of ribosomes [1,2]. Ribosomes are central in protein synthesis machinery and, therefore, ribosome content is a determinant of protein synthesis, and hence of cell growth and division [1–3]. Thus because of the high metabolic cost of rRNA synthesis, a tightly regulated transcription system is necessary to ensure that the mature rRNA level is coupled with cellular growth demands. In the yeast *Saccharomyces cerevisiae*, RNA pol I uses 60% of all ribonucleotides for its single gene target [1,4]. This extremely high transcription rate (TR) is possible due to the high density and speed of this RNA pol [2] and because of the large number of copies of the 35S rDNA gene. In most eukaryotes the rDNA gene exists as a long tandem repeat in one or in several chromosomes [5,6]. So, it would seem that the large TR required for rRNA synthesis needed during active cell growth and proliferation [7] cannot be achieved only by using RNA pol I at its maximum capacity, and multiplication of the rDNA gene is necessary. This is a slow form of TR regulation because the change in the rDNA copy number can be done only during genome replication using an unequal homologous recombination between sister chromatids [6,8,9]. rDNA loci are very dynamic genome regions whose copy number varies considerably in a single species between individuals, and even between cells in a single individual [5,10–13]. Apart from regulating the rDNA copy number, it is also possible to regulate rRNA synthesis by acting on RNA pol I transcription initiation [14] or elongation (see [15] for a recent review), and by controlling the proportion of active rDNA repeats. This last option is possible because repeats can exist as silenced chromatin covered by regularly packaged nucleosomal arrays or as a transcriptionally active and nucleosome-depleted copy [7,16]. It is likely that this mechanism is also slow as the change between chromatin states requires the passage of the replication fork through rRNA genes to reset nucleosome assembly [17].

In order to maintain its identity and physiological features, a cell should keep the concentrations of its molecules constant, or at least within a certain limited range. This homeostatic control involves maintaining the total amount of RNAs and proteins, which are the performers of genetic information. Total protein content is homeostatically controlled due to the high protein content of all cells and the high cost of protein synthesis (see [4] for a detailed discussion). This means that the ribosome concentration is strictly controlled as it is a key determinant of protein synthesis (see above) and, consequently, of the rRNA concentration ([rRNA]). [rRNA] depends on a dynamic equilibrium between its synthesis and decay rates. As stated above, the synthesis rate (SR) for rRNA is extremely high in actively dividing cells, which means that RNA pol I should be very efficiently controlled to adjust to cell requirements. The rate at which RNA pol I makes the rRNA precursor (also called 35-47S [5]) is called the nascent transcription rate (nTR, [18]) and accounts for the number of rRNA molecules made by a time unit. As the homeostatic equilibrium depends on concentrations, and not on numbers of molecules, changes in cell volume add another factor that should be taken into account. Change in cell volume is a phenomenon linked not just to progression along the cell cycle, but also to cell ageing as mother cells undergo progressive increases in size across generations [19], and also to the genotype of yeast strains because of the influence of many mutations on the cell cycle [20]. When cell volume changes, SR and nTR become numerically different because SR equals the nTR divided by cell volume [18]. However as the physical parameter that measures RNA pol activity is nTR, all kind of regulations act primarily on it.

We have previously shown that the nTR for RNA pol II is regulated differently in cells presenting symmetrical or asymmetrical cell division [21]. In that study we proposed three models or scenarios for nTR regulation. In scenario #1, symmetrically dividing cells increase the RNA pol II nTR in parallel with cell volume to keep SR constant. In this scenario RNA pol II is limiting and cell volume increase is accompanied by the volume-dependent recruitment of RNA pol II onto target genes, as recently demonstrated in *S. pombe* [22]. In scenario #3 however, the *S. cerevisiae* RNA pol II nTR remains constant in spite of cell volume changes by changing the RNA pol II cell concentration to avoid changes in cell physiology along successive asymmetric cell divisions [21]. In scenario #2, the *S. cerevisiae* RNA pol I nTR also remains constant with cell volume, but via a different mechanism: nTR regulation by limiting RNA pol I targets.

In this study, we extend the previous study by focusing on RNA pol I regulation in budding yeast as regards changes in cell volume. We have considered changes in cell volume during the cell cycle of an individual cell and when comparing the average cell volume in asynchronous actively growing yeast populations. By different types of experiments run to determine the RNA pol I nTR, we have found that budding yeast adapts the number of rDNA repeats to cell volume. We propose a model of regulation, based on that previously proposed by D. Shore [10] and T. Kobayashi [23,24] in which cell volume affects the activity of the Sir2 histone deacetylase which, in turn, regulates the homologous recombination at the rDNA locus and, thus, the possibility of varying the repeats copy number.

## Materials & Methods

### Yeast strains, media and growth conditions

The *S. cerevisiae* strains used herein are summarised in Supplementary Table 1. *S. cerevisiae* cells were grown in either liquid YPD (2% glucose, 2% peptone, 1% yeast extract) or YPGal (2% galactose, 2% peptone, 1% yeast extract) media. Experimental assays were performed with cells exponentially grown for at least seven generations until OD_600_ ~0.4-0.5 at 30°C.

The growth rate (GR) was calculated by growing 50 mL of yeast cultures in 250 mL flasks with shaking (190 rpm) at 30°C. Aliquots were taken every 60 min in the exponential phase and their OD_600_ were measured. GRs (h^−1^) were calculated from growth curves in YPD and/or YPGal and are shown in Supplementary Table 1.

For the experiment run to enlarge yeast cells, we used a strain with the only cyclin (*CLN3*) gene under the Gal1 promoter (BF305-15d, [25,26]). This strain only grows in galactose as a carbon source. To shut off the Gal1 promoter, an exponential phase culture in YPGal at 30°C (OD_600_ ~0.4) was collected, washed with pre-warmed YPD and resuspended at the same OD into YPD. Aliquots of cells were used to measure cell volume and number (see below), were then collected by centrifugation, the supernatant was discarded and pellets were frozen in liquid nitrogen. Three biological replicates of the experiment were done.

### *cln3* strain evolution experiment

For culture evolution purposes, *Δcln3* mutants were made from strain BY4741 by substituting the natural allele for the KanMX4 PCR-amplified cassette with the oligos described in Supplementary Table 2 following the lithium acetate protocol [27]. An rDNA copy number alteration due to the Li^+^ was discarded by qPCR (see below) in the obtained clones. Small colonies (1 mm diameter) were assumed to contain ~2×10^5^ cells and corresponded to progenies after 18 generations, starting from single ancestor cell [28]. The subcultures in liquid YPD medium were started and growth was followed by measuring OD_600_. Repeated sub-culturing was done by inoculating 10^4^ cells into 100 mL of fresh medium and allowing them to grow for 24 h to 5-6 OD_600_s (to about 10^8^ cells/mL, 12-13 generations). Each day an aliquot of the culture was spun down, the supernatant was discarded and the cell pellet was frozen onto liquid nitrogen. Four different colonies *Δcln3* (#9, 13, 20 & 24) were used.

### Elutriation

*S. cerevisiae* BY4741 cells were synchronised by elutriation by an Avanti^®^ J Series instrument from Beckman Coulter with a JE-5.0 elutriator rotor. The exponentially growing cells in YPD at 30°C (OD_600_ ~ 0.4) were placed inside the elutriation chamber at 4°C and cells were recovered by increasing the flow rate. Small cells were collected in cold YPD and keep on ice until 4×10^8^ cells were elutriated. Elutriation efficiency was confirmed by microscopy. Then cells were spun down at 4°C and resuspended in new pre-warmed YPD at 30°C. This was taken as time 0 for the experiment. The culture was incubated at 30°C and aliquots of 5×10^7^ cells were spun down every 8 min. The supernatant was discarded and frozen in liquid nitrogen. Three independent biological replicates of the whole experiment were done.

### Cell volume and flow cytometry

The median values of the cell volumes of the population were obtained by a Coulter-Counter Z series device (Beckman Coulter, USA), as previously described [21]. The absolute values (in fL) are shown in Supplementary Table 1. The percentage of cells to have initiated the S phase was determined by a flow cytometry analysis of propidium iodine-stained cells, as described in [29].

### RNA extraction, RT-qPCR for mRNA levels and mRNA half-life determination

To determine the amount of RNA, cells were grown in rich media until the exponential phase (see above). Then total RNA was extracted by phenol:chloroform extraction, as described in[30] in three biological replicates and was quantified by an OD_260_ estimation in a Nanodrop device (ThermoFisher Scientific) or using the Qubit RNA BR Assay kit (Thermo Fisher Scientific) measured in a Qubit fluorimeter for accurate quantification purposes.

The expression of *SIR2* and *UAF30* was measured by RT-qPCR in the DNase I-treated RNA samples extracted as previously described before being normalised against the *ACT1* mRNA levels. Specific primers were designed for this aim and are listed in Supplementary Table 2. The reverse transcription of mRNA was carried out using an oligo d(T)_15_VN with Maxima Reverse Transcriptase (Thermo Fisher Scientific). cDNA was labelled with SYBR Pre-mix Ex Taq (Tli RNase H Plus, from Takara) and the Cq values were obtained from the CFX96 TouchTM Real-Time PCR Detection System (BioRad).

### Determination of the nascent transcription and synthesis rates

For the total (RNA pol I + II + III) nascent transcription rates (nTR), a run-on experiment was used, followed by TCA precipitation onto a glass fibre filter. For these experiments, the *S. cerevisiae* cells were grown up to a DO_600_ 0.4-0.6. Three biological replicates were performed for each yeast strain. For each sample, two aliquots of 2.5×10^7^ cells were collected by centrifugation at 4,000 rpm for 3 min in a 1.5 mL Eppendorf tube. Pellets were resuspended in 0.5 mL of 0.5% sarkosyl and spun down again by centrifugation under the same conditions as above. Supernatants were discarded to completely eliminate sarkosyl. Then the pellet was resuspended in 7.2 μL of distilled water (for the total nTR samples) or in 7.2 μL of an α-amanitin solution (100 μg/mL) (for RNA pol I + III nTR samples). The run-on pulse was performed by adding to the cell pellet, per sample, 9.87 μL of the transcription mix, composed of 7.5 μL of 2.5× transcription buffer (50 mM Tris-HCl pH 7.7, 500 mM KCl, 80 mM MgCl2); 1 μL of the rNTP mix (10 mM each ATP, CTP and GTP); 0.375 μL of 0.1 M DTT and 1 μL of UTP (0.9 μL of 3 μM cold UTP, plus 0.1 μL of 3 μM [α-^33^P] UTP, Perkin Elmer, 3000 Ci /mmol, 10 μCi/μL), at a final volume of 18.75 μL. To allow transcription elongation, the mix was incubated with agitation (650 rpm) for 5 min at 30°C. The reaction was stopped by adding 82 μL of cold distilled water to the mix and leaving it on ice. To measure the total amount of radioactivity present in the mix (‘Total’), 15 μL of the reaction were directly spotted onto glass fibre paper discs and dried in an aerated heater at 65°C. Another 15 μL aliquot of the mix was spotted onto glass fibre discs and precipitated by soaking it in 4 mL of 10% (v/v) of trichloroacetic acid (TCA) at 4°C for 20 min. TCA was removed and a new wash of 4 mL of cold TCA (10% v/v) was added for 10 min. TCA was removed and discs were washed with 3 mL of cold 70% (v/v) EtOH. Discs were dried again in a heater at 65°C. Once dried, 5 mL of a scintillation cocktail (Normascint #22, Scharlau) were added to each disc for radioactive counting. Each of the three biological replicates was performed in three technical replicates for TCA and two for total and averaged. For each individual sample, the incorporation was calculated as “TCA/Total”. Synthesis rates (SR) were estimated by dividing the nTR values by cell volume (see above).

### Chromatin immunoprecipitation of RNA pol I

The recruitment of RNA polymerase I to chromatin was assayed by a chromatin immunoprecipitation (ChIP) analysis with a specific antibody for RNA pol I. Cells were grown in rich medium until the exponential phase. Crosslinking between proteins and the associated DNA was performed by adding formaldehyde to 0.75% for 15 minutes. Samples were sonicated 12 times (30 sec on, 30 sec off) at the high intensity in a Bioruptor (Diagenode) device. Chromatin fragments were immunoprecipitated using an antibody against RNA polymerase I subunit 135 (A135) coupled to magnetic beads. DNA fragments were then amplified by RT-qPCR and the Cq values were analysed. SYBR Pre-mix Ex Taq (Takara) was used for RT-qPCR following the manufacturer’s instructions and the reaction was performed in the CFX96 TouchTM Real-Time PCR Detection System (BioRad). The primers designed to detect ribosomal genes (25S, 18S) are listed in Supplementary Table 2. Their localisation on the rDNA repeat is shown in Figure 3D.

### qPCR quantification of rDNA repeats

The estimation of the number of ribosomal RNA gene (rDNA) repeats was measured by RT-qPCR and normalised against the *ACT1* gene signal. For this purpose, genomic DNA was purified by a standard phenol:chloroform extraction [31] and precipitated with ethanol. Primers were designed against a specific region in the rDNA cluster (5.8S) and are listed in Supplementary Table 2. DNA was labelled with SYBR Pre-mix Ex Taq (Tli RNase H Plus) from Takara and the Cq values were obtained from the CFX96 TouchTM Real-Time PCR Detection System (BioRad).

### Western blot analysis

Protein extraction from 1-2×10^8^ yeast cells was done by suspending the cell pellet in 200 μL of NaOH 0.2M. After 5 minutes, samples were centrifuged and then mixed with 100 μL 2X SDS-PAGE loading buffer. Then they were incubated at 95°C for 5 minutes to completely extract proteins from cells. For the Western blot analysis [32] similar protein amounts were injected into SDS-PAGE acrylamide gels under reducing and denaturing conditions. Protein antibodies against A190 and A135 RNA pol I subunits were assayed in this study and were provided from O. Calvo & C. Fernández-Tornero. Glucose 6-phosphate dehydrogenase (G-6-PDH) was used as an internal reference by re-incubating the same blots with a specific antibody (22C5D8 from Thermo Fischer Scientific).

## Results

### The total nascent transcription rate (nTR) remains constant along the cell cycle in yeast

In a previous work, we showed that RNA pol II nTR remains constant while increasing cell volume during the cell cycle in *S. cerevisiae.* So we then wondered what would happen to the other nuclear RNA polymerases. To find an answer to this question, we performed a synchronisation of yeast cells by elutriation. We obtained a sample of newborn daughter cells at 4°C. This sample was allowed to grow by placing cells at 30°C for 50 min. The cell number was constant (Figure S1A), and an average increase in size of about 28% for up to 50 min took place (Figure S1B) when cells started budding (Figure S1C). We then determined the total nTR of each sample by run-on incorporation. We used a simplified protocol with TCA precipitation of the labelled nascent RNA into glass fibre filter discs (filter run-on: see M&M). Given that RNA pol II only accounts for 25% of the total nTR [1] and remains constant along the cell cycle [21] this result informs about RNA pol I+III nTR behaviour. Figure 1A shows that total nTR is constant in spite of a significant increase in cell volume. This means that the effective RNA synthesis rate considering [rRNA] concentration (SR) (see the Introduction) lowers with cell volume (Figure S1D). We conclude that the nTR for the sum of RNA pol I, which accounts for 60% of the total nTR [1], and RNA pol III (15%) were constant.

**Figure 1.**
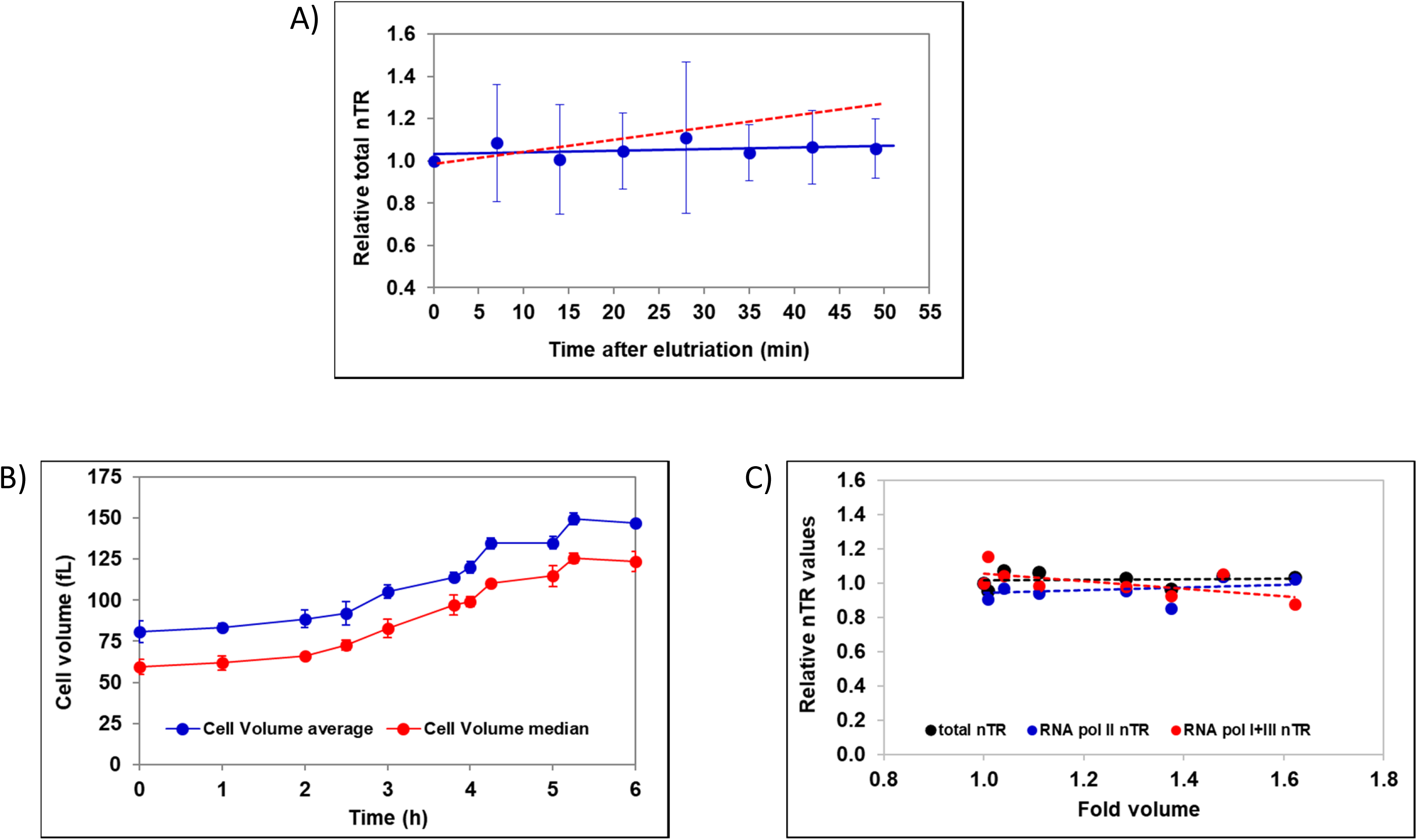
The total nascent transcription rate (nTR) is constant in *S. cerevisiae* along the cell cycle and cell size increase. A) Cells were synchronised by isolating newborn small cells by elutriation. After resuming growth, cell aliquots were taken and used for nTR determination by filter run-on. The nTR profile vs. time is shown. The total RNA synthesis rate (SR) was calculated by dividing the nTR by cell volume and represented vs. the relative cell volume (see Figure S1B). The red line shows the expected results if the nTR had increased in parallel to cell volume. B) We induced cell enlargement by shutting off the *CLN3* gene in a strain in which Cln3 was the only cyclin and it was transcribed from the *GAL1* promoter. Cells were grown in YPGal and then shifted to YPD. At the indicated times, aliquots were taken, and cell growth (see Figure S2) and volume were measured. C) The nTR was determined by filter run-on in the absence (the total nTR: I+II+III) or presence (RNA pol I+III nTR) of α-amanitin. The RNA pol II nTR was calculated as the difference between the total and α-amanitin values. Data are relative to the 0 time point. Data represent the average and standard deviations from three biological replicates of the experiment. Note that the SDs in graph C) are too small to see error bars.

This result differs from that found by other authors who have worked in cells with symmetric division. In both human fibroblasts [33] and *Schizosaccharomyces pombe* [34] RNA pol I nTR was found to increase in parallel to cell volume. In the latter case, P. Nurse & collaborators used temperature-sensitive *S. pombe* mutants to stop the cell cycle and provoke a marked increase in cell volume. We decided to use a similar approach in *S. cerevisiae* by using a mutant strain with only one cyclin gene, *CLN3*, under the control of the Gal1 promoter [25]. When changing it from galactose (YPGal) to a glucose-based medium (YPD), cells stopped the cell cycle at G1-S and started enlarging cell volume from 1h onwards (Figure 1B), which was the main cause of the OD_600_ increase (Figure S2A) because the cell number remained constant from 2 h of incubation (Figure S2B). We determined the range of time of the samples at which the average cell volume grew almost linearly with time up to about 200% as regards the original 65 fL size (Figure 1B). Then we performed filter run-on with those cell samples in duplicates with and without α-amanitin, a potent inhibitor of RNA pol II [35], which allowed us to discriminate the nTR from RNA pol II and RNA pol I+III. Both similarly showed a flat profile (Figure 1C), which demonstrates that even if a large volume increase is caused by the growth of a daughter cell along G1, *S. cerevisiae* keeps RNA pol I constant and behaves differently from symmetrically dividing *S. pombe*. The minor contribution of RNA pol III to α-amanitin-resistant transcription made it difficult to assess if it was also constant.

### The nascent transcription rate of RNA pol I is proportional to the ploidy of yeast strains and varies with cell volume

In a previous study [21], we used non-synchronised exponentially growing cultures of yeast strains with different average cell volumes. The set of strains included the wild-type haploid strain BY4741, two haploid mutants, *cln3* and *whi5*, respectively with large and small volumes, and three polyploid (2n, 3n & 4n) strains created by D. Pellman’s group [36] (with cell sizes of approximately 2×, 3X and 4X as regards BY4741). All the strains grow similarly (Supplementary Table 1), which excludes any difference in the TR due to changes in growth rates [37]. We found that the total nTR increased linearly with volume in polyploid cells because of the parallel increase in genome copies (the nTR/genome remained constant), and we postulated a scenario for RNA pol I regulation in which the nTR of this RNA pol strictly depended on the abundance of its substrate: 35S rDNA [21]. Then we analysed the behaviour of haploid mutant strains *whi5* and *cln3* by filter run-on, and by chromatin immunoprecipitation of the second largest RNA pol I subunit: A135. Figure 2A-B depicts how the total nTR per genome copy is much higher in *cln3* and slightly lower in *whi5* than in the wild-type and the diploid strains. Thus as differences in the nTR cannot be caused by changes in ploidy for haploid mutants with a different cell size, we hypothesised that the variation in the RNA pol I substrate could be due to a change in the number of rDNA repeats.

**Figure 2.**
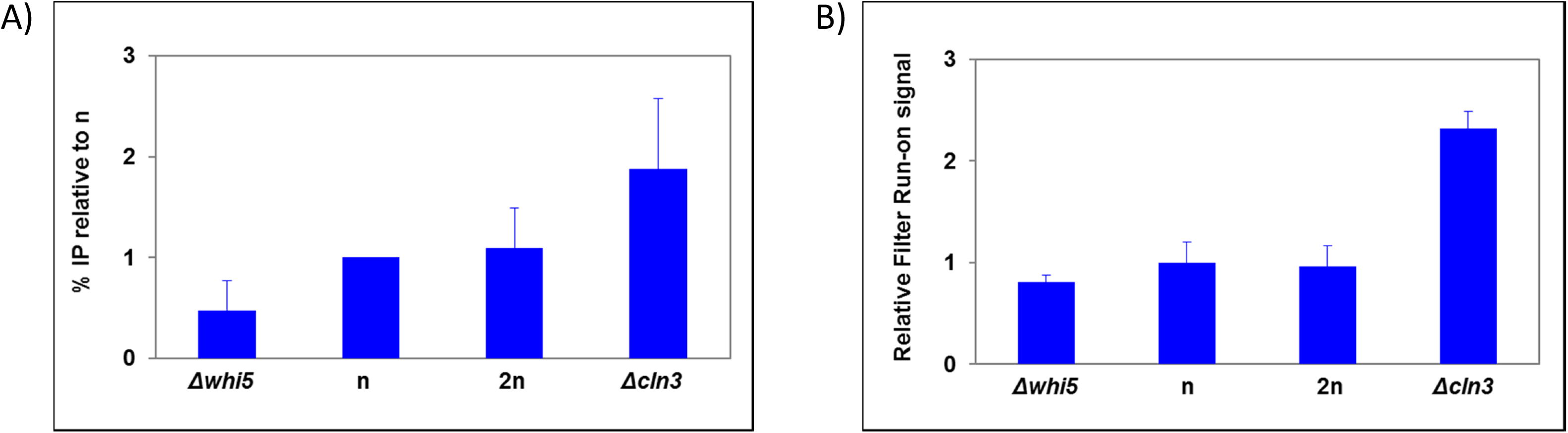
The RNA pol I nTR per genome copy changes in haploid cell size mutants. The nTR was determined in a series of haploid yeast strains with different cell volume (see Supplementary Table 1) and a diploid (2n) control strain for the RNA pol I by chromatin immunoprecipitation of its second largest subunit (A135) (A) or for the total nTR by filter run-on (B).

**Figure 3.**
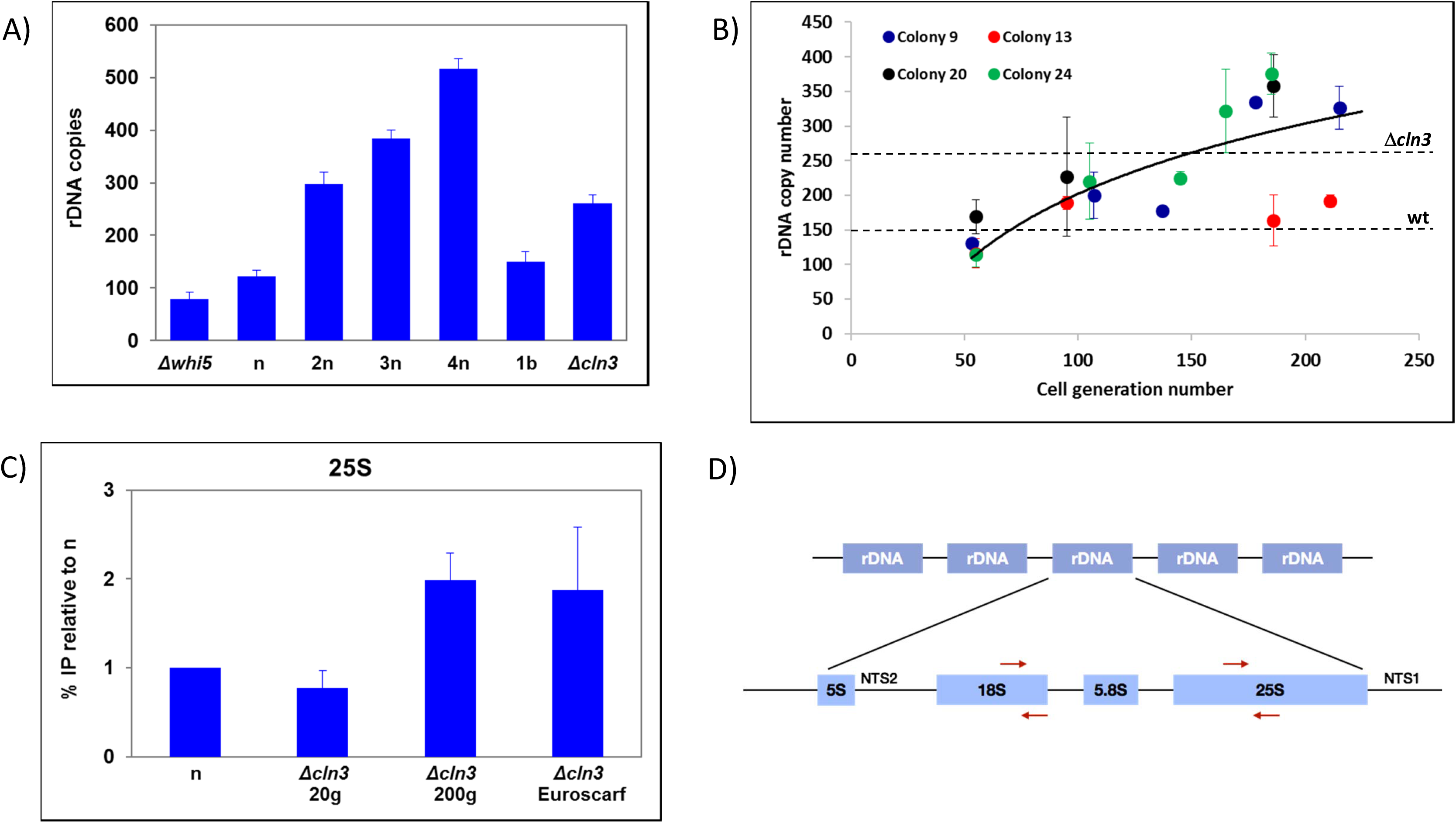
The number of rDNA repeats evolves in a *cln3* mutant to match the optimum value of the RNA pol I nTR required for its cell volume. A) The number of rDNA repeats varies in different strains in a way that depends on ploidy, except for cell size haploid mutants (see Figure S3). B) In a *cln3* mutant, the copy number is small, similar to its parental wt haploid (n: BY4741) when it is a recent transformant, but increases along with the number of generations for four independent clones (#9,13,20,24). The dashed horizontal lines mark the repeat copy number for the Euroscarf *Δcln3* (BQS2006) and wt (BY4741) strains. C) The RNA pol I nTR of the *cln3* mutants varies according to the rDNA repeat number, as determined by chromatin immunoprecipitation of the second largest RNA pol I subunit (A135) onto either the 25S (this figure) or 18S (Figure S3C) transcribed regions. *Δcln3* Euroscarf is BQS2006 strain (see Supplementary Table 1) D) A scheme of the rDNA locus and the probes used for qPCR is shown. Strain NOY408-1b (“1b”, see Supplementary Table 1), with a known rDNA copy number (150 repeats, see [28]), was used as an internal control.

### The rDNA copy number increases in a *cln3* mutant to compensate an increased cell volume

The rDNA gene in *S. cerevisiae* forms tandem repeats between 100-200 copies [9]. The repeat copy number has been shown to vary between different mutants [38]. Therefore, we hypothesised that the number of rDNA repeats increased in the *cln3* mutant and decreased in *whi5* to adjust its number to the average cell size. To test this, we measured the rDNA repeat number in the series of yeast strains by qPCR. Figures 3A and S3A show that the total copy number in relation to a single copy gene control (*ACT1*) is more than double in the *cln3* mutant and is slightly lower in the *whi5* mutant compared to BY4741. In fact the relative rDNA change in both mutants as regards BY4741 (2.2 and 0.9) comes very close to the relative changes in the observed RNA pol I nTR (data from Filter run-on GRO and ChIP with A135 in Figure 2).

If the mutants had a different repeat copy number from the wild-type strain that they derive from, we hypothesised that it would be due to a secondary effect after deleting the target gene. This is because the cell, where deletion occurs, has a wild-type number of rDNA repeats. We hypothesised that the rDNA copy number would be adjusted after gene deletion by successive steps across genome replications [28]. To test this, we did a 200-generation culture of four freshly obtained *Δcln3* mutants (<20 generations). We first checked the cell size of these mutants and found that it was indistinguishable from an old *cln3* mutant obtained from the Euroscarf collection in which we performed previous experiments: 83±10 fL (Figure S3C, Supplementary Table 1). We also checked that these four mutants in colonies of less than 20 generations had a wild-type rDNA copy number. Then we cultured these freshly made *Δcln3* mutants in YPD for approximately 200 generations by refreshing cultures daily in new medium every 10-12 generations. Figure 3B illustrates that all the *cln3* strains progressively increase the number of rDNA repeats to up 300 on average, although variability among strains was wide. Thus it would seem that a fresh *S. cerevisiae cln3* mutant had an intrinsic enlarged size because of its longer G1 period [26,38] and not as a result of an altered rDNA copy number. Its larger size probably caused a defect in the effective synthesis rate (SR) of 35S rRNA that was ultimately compensated by the plasticity of the rDNA locus to increase the repeat copy number and, consequently, nascent transcription. In fact the measurements of elongating RNA pol I by immunoprecipitation with Anti-A135 Ab confirmed that the actual RNA pol I transcription increased in the newly generated *Δcln3* cells across generations (Figure 3C and S3B).

### A model for RNA pol I transcription regulation by cell volume

We previously published a model for the regulation of RNA pol I transcription with cell volume, which is based on a large excess and a high affinity of this RNA pol to its targets (scenario #2 in [21]). This model predicts that [RNA pol I] does not decrease with cell volume to keep the nTR constant (differently from RNA pol II, see [21]). We have used the set of yeast strains with different cell volumes (see Figure S3C) for the Western quantification of the two main subunits of this RNA pol to find that the RNA pol I concentration remained constant (Figure 4).

**Figure 4.**
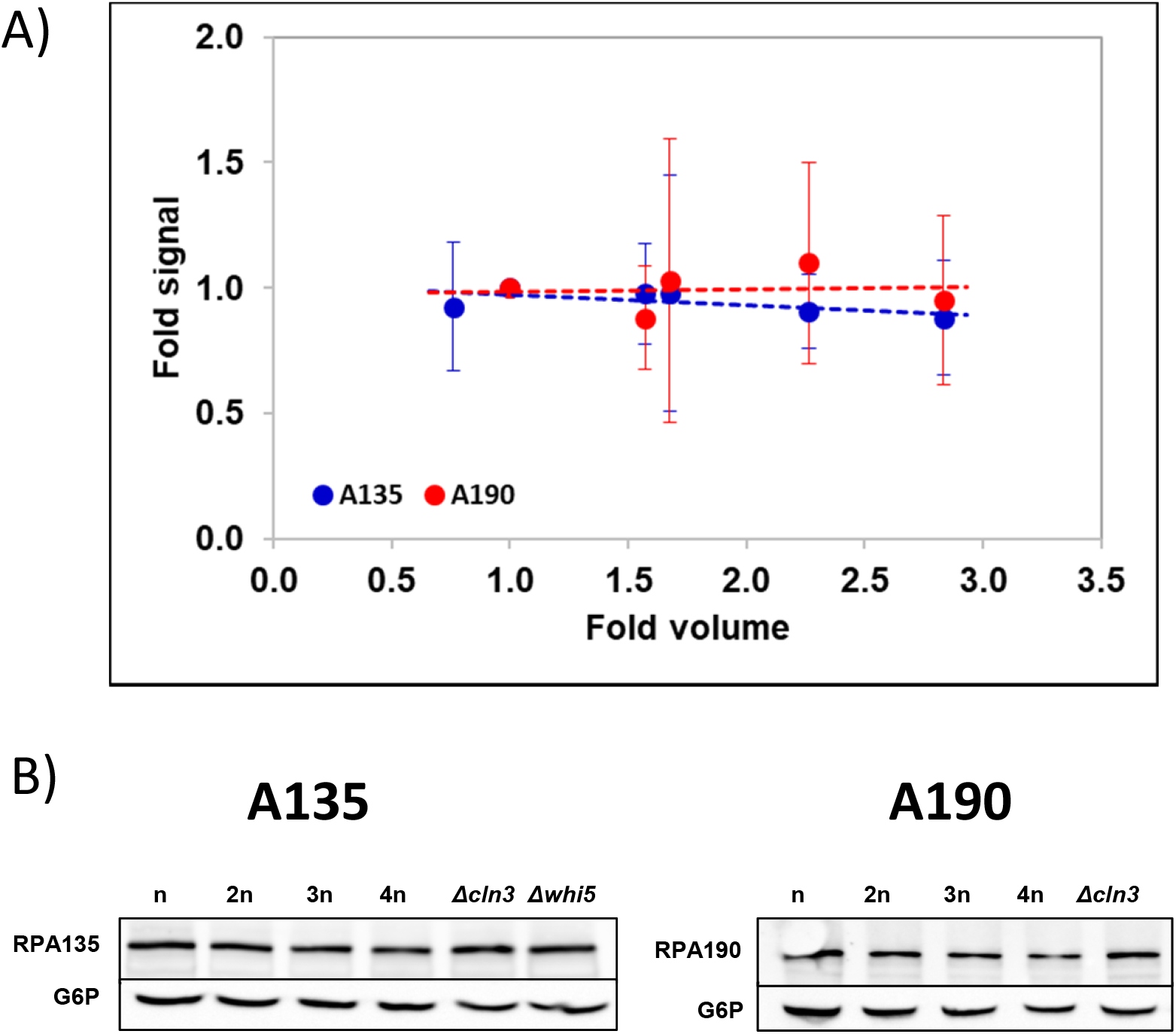
The cellular concentration of RNA pol I does not vary with cell volume. A) The RNA pol I concentration in different polyploids (n-4n) and haploid mutants (*whi5*, *cln3*) shown in Figure 3A was calculated by Western blot using approximately the same amount of total protein and correcting by G6P-DH signal. An antibody against the second largest RNA pol I subunit (A135, blue dots) and one against the largest RNA pol I subunit (A190, red dots) are shown. The graphs represent the average and standard deviations from three biological replicates. B) In both cases, an example of a Western blot is shown below.

In scenario #2 of [21] the total nTR was thus driven by the actual number of rDNA gene copies present in each strain. For the previously described 200-generation-evolved *cln3* mutants, this would appear to be modulated by an amplification of the rDNA repeats occurring along many genome replications. T. Kobayashi’s group has shown that the rDNA locus is very plastic and undergoes expansions and contractions by unequal sister-chromatid recombination, which produces two cells with different repeat copy numbers, followed by a gradual evolution (see [28,40]). This pathway fits our previous results with *cln3* mutants. A recent paper [23], postulates that the control of homologous recombination is due to the action of histone deacetylase Sir2 which, in turn, is controlled at the transcription initiation level by the upstream activator factor (UAF) complex. The UAF is both an activator of RNA pol I transcription and a repressor of *SIR2* transcription by RNA pol II. Here we adapt this model to account for our observations as regards cell volume. Our model (Figure 5A) assumes that both the UAF and RNA pol I have very high affinity constants for the rDNA promoter. Under steady-state growth conditions, both proteins saturate the available (open chromatin) repeat copies. Under steady conditions the UAF is in a limiting amount [23] whereas RNA pol I is in excess [7,21]. When the *CLN3* gene is deleted after transformation, the wild-type cell instantaneously becomes a *Δcln3* mutant and changes its cell cycle to become larger than the wild type. As most proteins maintain their concentration independently of cell volume, the number of the UAF and RNA pol I molecules/cell increases proportionally to cell volume. This has no effect on RNA pol I behaviour because the large molar excess over its targets renders the change in the number of free molecules irrelevant. For the UAF, the consequence is different because it provokes a change in the number of free molecules proportional to the volume increase and then, because of the constant rDNA copy number, in the free [UAF]. UAF has also a low affinity for the *SIR2* promoter, although much lower than for rDNA [23]. Therefore, an increased concentration of free UAF leads to the repression of *SIR2* gene. Thus in larger cells, the UAF will repress *SIR2* transcription which, in turn, will cause the de-repression of the homologous recombination and promote copy number increase [28]. When the number of rDNA repeats reaches the necessary number to sequester most of the free UAF molecules present in the new cell volume, the system will return to the stationary state.

**Figure 5.**
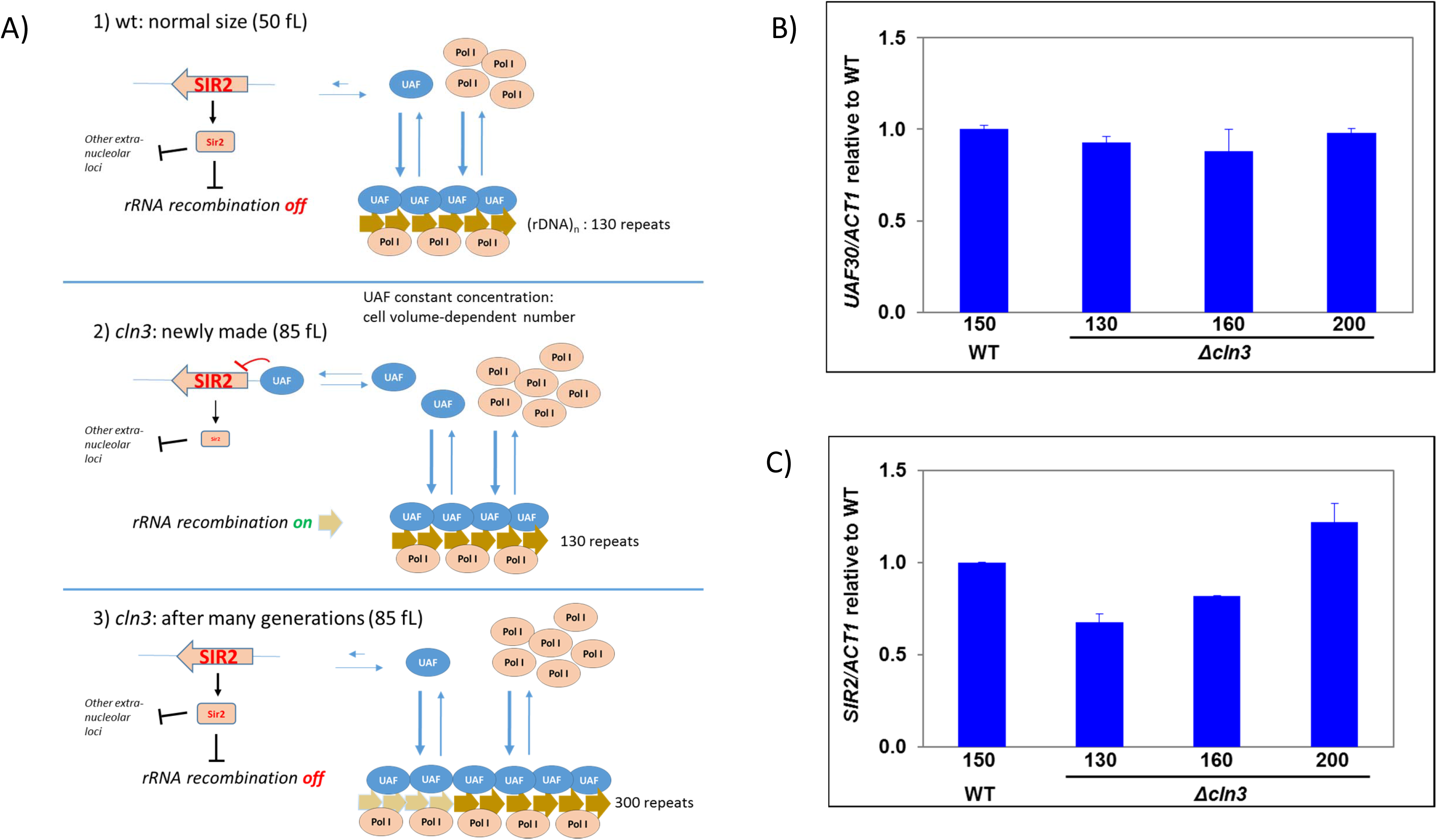
Model for rDNA copy number regulation by cell volume. A) A model for cell size control on the rDNA repeat copy number is shown (see the main text for details). The relative levels of *UAF30* (B) and *SIR2* (C) to a reference gene (*ACT1*) in the *cln3* strains of different numbers of generations during the evolution experiment determined by RTqPCR are shown.

We study this homeostatic control of the rDNA repeat copy number using a differential equation-based mathematical model (Appendix in Supplementary Information). Intriguingly, our mathematical model corresponds to an integral feedback mechanism to control the concentration of rDNA repeats at a prescribed set point. Such integral feedbacks have recently been shown to be essential for robust perfect adaptation when faced with environmental perturbations [41]. We refer readers to the Supplementary Information for details on the mathematical modelling.

A prediction of this model is that the total cell [UAF] will be always constant, but excess free [UAF] will repress *SIR2* expression in freshly made *cln3* mutants and will go back to its wild-type levels after 200 generations. Figure 5B shows that *UAF30* subunit mRNA expression is constant in the *cln3* strains with a different rDNA copy number, but *SIR2* expression is lower in the *cln3* strains with small rDNA copy number and increases to wild-type levels in the strains with a larger rDNA copy number (Figure 5C).

## Discussion

Our previous study [21] found that in cells with asymmetric cell division, such as budding yeast, the size difference between mother and daughter cells imposes a restriction to the way eukaryotic cells deal with the problem of controlling the total RNA pol II TR as regards cell volume. The conceptual problem arises when we consider that the actual number of transcripts produced by a time unit (nTR) strictly depends on the activity of RNA polymerases in a fixed number of gene templates. The actual SR in chemical kinetic terms involves the concentration of transcripts, instead of their number, which means that cell volume changes become a new player in the equilibrium (see [18]). In fact the real objective of regulation is the concentration of transcripts itself, which can also be controlled at the decay (degradation + dilution) level on the other side of the equilibrium reaction to achieve homeostasis [4].

How the cell controls the activity of its RNA polymerases (i.e. their nTR) to get the right SR depends on the symmetry of cell division. Symmetrically dividing cells, such as human fibroblasts and *S. pombe*, increase the number of elongating RNA pol II by keeping its cellular concentration constant independently on cell volume, while maintaining a very strong association constant with gene promoters [33–34]. In this way the equilibrium is strongly displaced towards the chromatin binding of RNA pol II, which assures that the increase in RNA pol II molecules with cell volume, required to maintain the concentration constant, will directly increase the nTR. We called this “scenario #1” in our previous study [21]. It has been recently demonstrated in *S. pombe* that RNA pol II amount scales with cell volume during the cell cycle and has a direct consequence on transcription initiation rates [22]. Due to the asymmetric division in budding yeast, nTR cannot increase with cell volume because it would provoke a never-ending increase in daughter cells every generation [21]. Instead we found that *S. cerevisiae* lowers the SR through volume increase because the RNA pol II concentration is down-regulated by cell volume and mRNA homeostasis is maintained by a compensatory increase in global mRNA stability [21].

The conceptual problem solved for RNA pol II in different organisms is applicable to the other nuclear RNA pol. Our previous study suggested that in budding yeast, RNA pol I also has a constant nTR due to an excess amount of this RNA pol regarding its rDNA template (which is limiting). We proposed a new scenario (#2), in which the rise in the nTR in large-sized polyploid cells would be caused by their increase in ploidy, which proportionally augments the total number of rDNA repeats. However, we found that other yeast strains with smaller or bigger sizes than the wild type, but with haploid genomes (e.g. *cln3* and *whi5*), also adapt their total nTR to their cell size. We suggested that it could be related with the plasticity of rDNA repeats, which are known to be variable in number [38]. In the present study, we performed a series of experiments that confirmed our hypotheses. We also found that the number of rDNA repeats in budding yeast adjusted to the required RNA pol I SR by a mechanism based on transcriptional control of the histone deacetylase *SIR2* gene (Figure 5). Our model of rDNA copy number control by cell volume is based on T. Kobayashi’s previous model [23,24] called “Musical chair”. This model predicts that, similarly to the scenarios proposed for RNA pol II transcription rate control (see above), a limiting amount of a factor (UAF), together with its high binding constant to the rDNA promoter, would lead the concentration of free UAF to be negligible in wild-type cells, but exquisitely dependent on the number of rDNA repeats. Iida and Kobayashi [23] demonstrated that the UAF is also able to bind the *SIR2* gene promoter, albeit with a lower binding constant than for an rDNA promoter, by acting as a repressor. In this way Sir2 histone deacetylase activity becomes dependent on the number of free UAF molecules. The silencing activity of Sir2 on an ncRNA gene placed within the rDNA repeat is necessary to repress the homologous recombination between rDNA repeats during replication, which is the origin of repeat copy number variability [6,23,24]. This variability would then allow the evolution of the appropriate rDNA copy number for its genetic features and cell volume.

We hypothesised that any change in cell volume with no change in rDNA repeats would alter the concentration of free UAF molecules that repress *SIR2* expression. Then we found that the increase in rDNA copies occurring in a freshly made *cln3* mutant (Figure 3B), with a wild-type copy number, but a much larger cell size (Figure S3C), provoked a decreased *SIR2* expression, which was recovered to reach the wild-type values (Figure 5B) along the progressive increase in rDNA repeats seen across successive generations (Figure 3B). This change occurred without any changes in UAF30 expression (Figure 5C).

According to the model, RNA pol I concentration is always in excess and does not need to change (Figure 4), unlike what we found for RNA pol II [21]. Why is there a different solution for an identical problem in RNA pol I and II? We think that the differences between the 35S and RNA pol II genes have conditioned the evolution of different regulatory mechanisms for both RNA polymerases. rRNA transcription should be able to reach much stronger levels than that of any of the RNA pol II genes given the need for huge numbers of ribosomes during active growth [1,3,4,7]. Eukaryotic cells have solved this by evolving a specialised polymerase that has a single gene template with many repeated copies [2]. The repeated nature of the rDNA locus is prone to cause a homologous recombination [24]. Direct repeats involve always a risk of deletion and genome re-arrangements [6,8]. In this case, however, they bring the opportunity to alter the rDNA copy number and, thus, the total nTR without changing the nTR/copy. In this way RNA pol I can be controlled in the short term at the transcription initiation level (similarly to RNA pol II genes). Moreover in the long term, the RNA pol I nTR can be changed by altering the chromatin structure of rDNA repeats [7,42] or by changing its copy number during genome replication. This difference with RNA pol II has probably provoked evolution to select a different way to deal with the cell volume problem in the RNA pol I TR. The low stability of RNA pol II transcripts, compared to the rRNA one, is another aspect that could explain the difference in regulatory strategies for RNA pol II and I regarding cell volume. mRNAs can compensate SR changes by modifying their stability [21] but very stable rRNA cannot. An interesting question that arises here is what happens to RNA pol III (the remaining nuclear eukaryotic RNA polymerase). This RNA pol transcribes an intermediate number of genes (about 400 in budding yeast). Approximately two thirds of them are tRNAs and one third is rRNA 5S repeats. tRNAs are variable in copy number (up to several dozens), but do not usually vary in a given species. Conversely, 5S genes are much more repeated, and its number is within the range of the longer rRNA gene transcribed by RNA pol I [43]. Interestingly, 5S genes form part of the rDNA repeats in the *Sacccharomycotina* clade [44], which comprises asymmetrically dividing yeasts. However in other yeasts and most other eukaryotes with symmetric cell division, 5S genes are usually dispersed along the genome [45].

Finally, another question to arise from our model is the possible additional effects that the change in Sir2 activity can have on other cellular processes. This histone deacetylase is responsible for maintaining chromatin silencing at telomeric regions, and for mating type loci and rDNA [40]. At rDNA, Sir2 acts in a complex called RENT, which interacts with the 35S promoter. Although Sir2 seems dispensable for RNA pol I transcription, its tethering to the 35S promoter seems to be important to strike a balance between Sir2 action in rDNA and at telomeres (see [40] for a detailed description). Any shrinkage in the rDNA array releases Sir2 and causes both an increase in telomeric and mating-type gene silencing and a proportional down-regulation of the *SIR2* gene to repeat loss [10]. The down-regulation of Sir2 can be explained by the UAF repression described in the Musical Chair model [23]. D. Shore’s group proposed that there is equilibrium between the nucleolar and non-nucleolar pools of the Sir2 protein, and that rDNA “buffers” the amount of Sir2 in a cell [10]. Quite unexpectedly, yeast strains with fewer rDNA repeats have normal growth rates [10,28]. We have also found that fresh *cln3* mutants with a small rDNA copy number have no growth defects (not shown). The defects in rDNA silencing by Sir2 affect DNA replication and promote increased ARS firing and DNA recombination, which lead to genome instability and lethality [46]. The decreased Sir2 activity in the *cln3* low repeat copy number may cause increased cell death. This has been proposed by A. Amon [47] who showed that ERC accumulation provokes fast ageing and death even in young cells. We found that, indeed, fresh *cln3* strains show a death rate that doubles either the wild type or older *cln3*, with an increased rDNA copy number (not shown). The *cln3* evolution mechanism from 130 to 300 rDNA repeats is an interesting topic to be investigated.

## Acknowledgements

We thank D. Pellman, M. Aldea and T. Kobayashi for the yeast strains, and O. Calvo and C. Fernández-Tornero for the anti-RNA pol I antibodies. We thank D.A. Medina, A. Arco and M.C. Centeno for collaborating in the initial experiments of this work. We also thank J. Pla for generously helping with the elutriation experiment. We are grateful to P. Alepuz, F. Navarro, M. Aldea for their helpful discussion. This work has been supported by grants from the Spanish Ministry of Economy and Competitiveness, and European Union funds (FEDER) [BFU2016-77728-C3-1-P to S. C.], [BFU2016-77728-C3-3-P and BFU2015-71978-REDT to J.E.P-O], and from the Regional Valencian Government [PROMETEO II 2015/006 & AICO2019/088 to J.E.P-O]. M. Barba-Aliaga was funded by a pre-doctoral research grant (FPU17/03542) from the Spanish Ministry of Science, Innovation and Universities. Funding for open access charge: [AICO2019/088].

---

**Figure S1.**
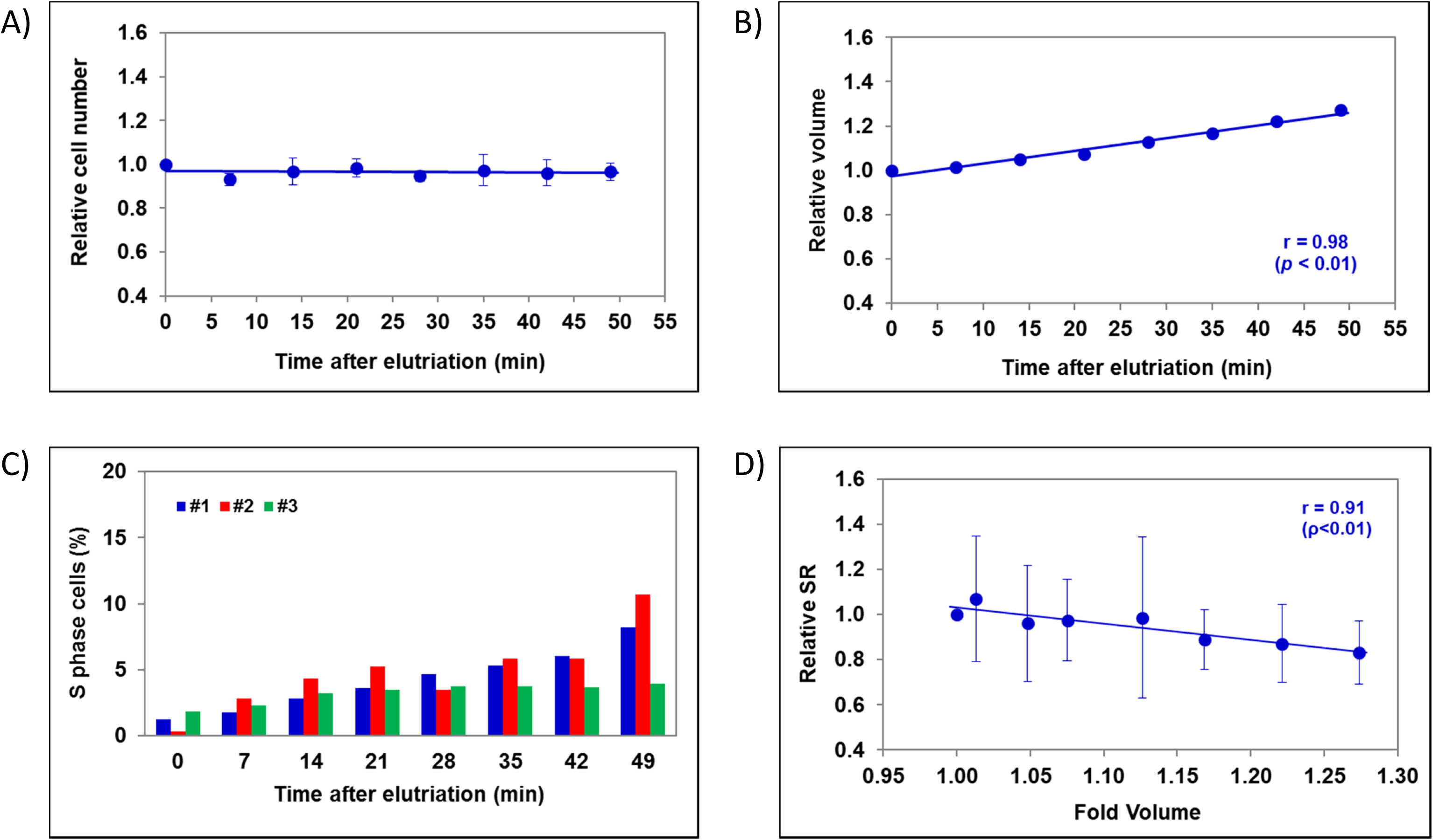
The total synthesis rate (SR) decreases in *S. cerevisiae* along the cell cycle. Cells were synchronised by isolating newborn small cells by elutriation. After resuming growth, cell aliquots were taken and used for cell number (A) and cell volume (B) determinations by a Coulter Counter device. The percentage of S phase cells was determined by flow cytometry (C). In (D), the total RNA synthesis rate (SR) was calculated by dividing the nTR (Figure 1A) by cell volume, and represented vs. the relative cell volume. Data represent the average and standard deviations from three biological replicates of the experiment. Note that the SDs in graph B) are too small to see error bars. This figure is complementary to Figure 1A.

**Figure S2.**
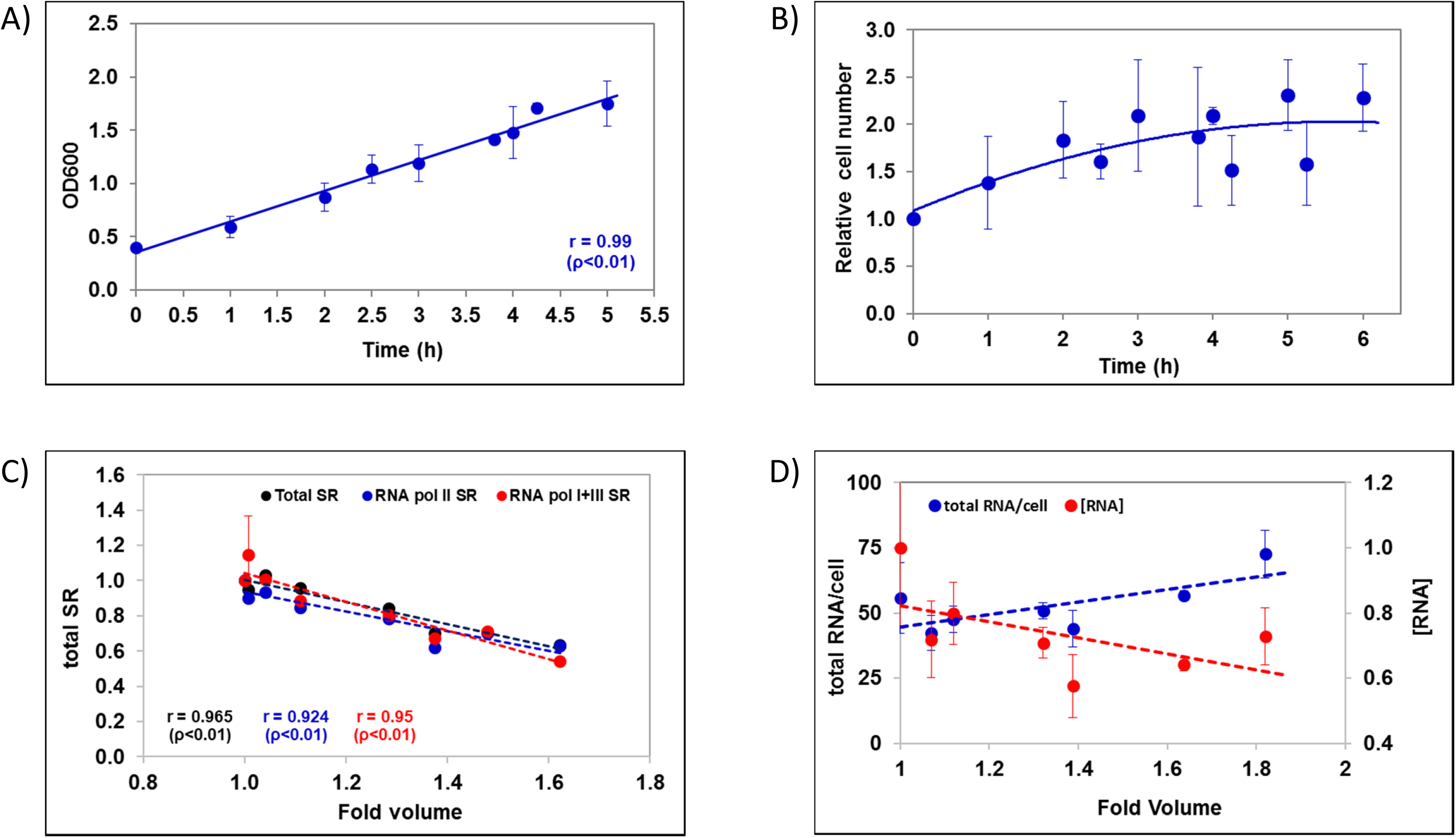
Increase in cell size in the G1/S blocked cells in *S. cerevisiae* is accompanied by SR decrease. We induced cell enlargement by shutting off the *CLN3* gene in a strain in which Cln3 was the only cyclin and it transcribed from the *GAL1* promoter. Cells were grown in YPGal and then shifted to YPD. At the indicated times, aliquots were taken and cell growth by OD_600_ (A) and cell number (B) were measured. The nTR was determined by filter run-on in the absence (total nTR: I+II+III) or presence (RNA pol I+III nTR) of α-amanitin (see Figure 1C). Then the synthesis rate (SR) was calculated by dividing the nTR by the relative fold volume. The RNA pol II SR was calculated as the difference between the total and α-amanitin values. Data are relative to the 0 time point (C). Finally, total RNA amount per cell was calculated by extracting RNA from a known number of cells (see M&M). Data represent the average and standard deviations from three biological replicates of the experiment. This figure is complementary to Figure 1B-C.

**Figure S3.**
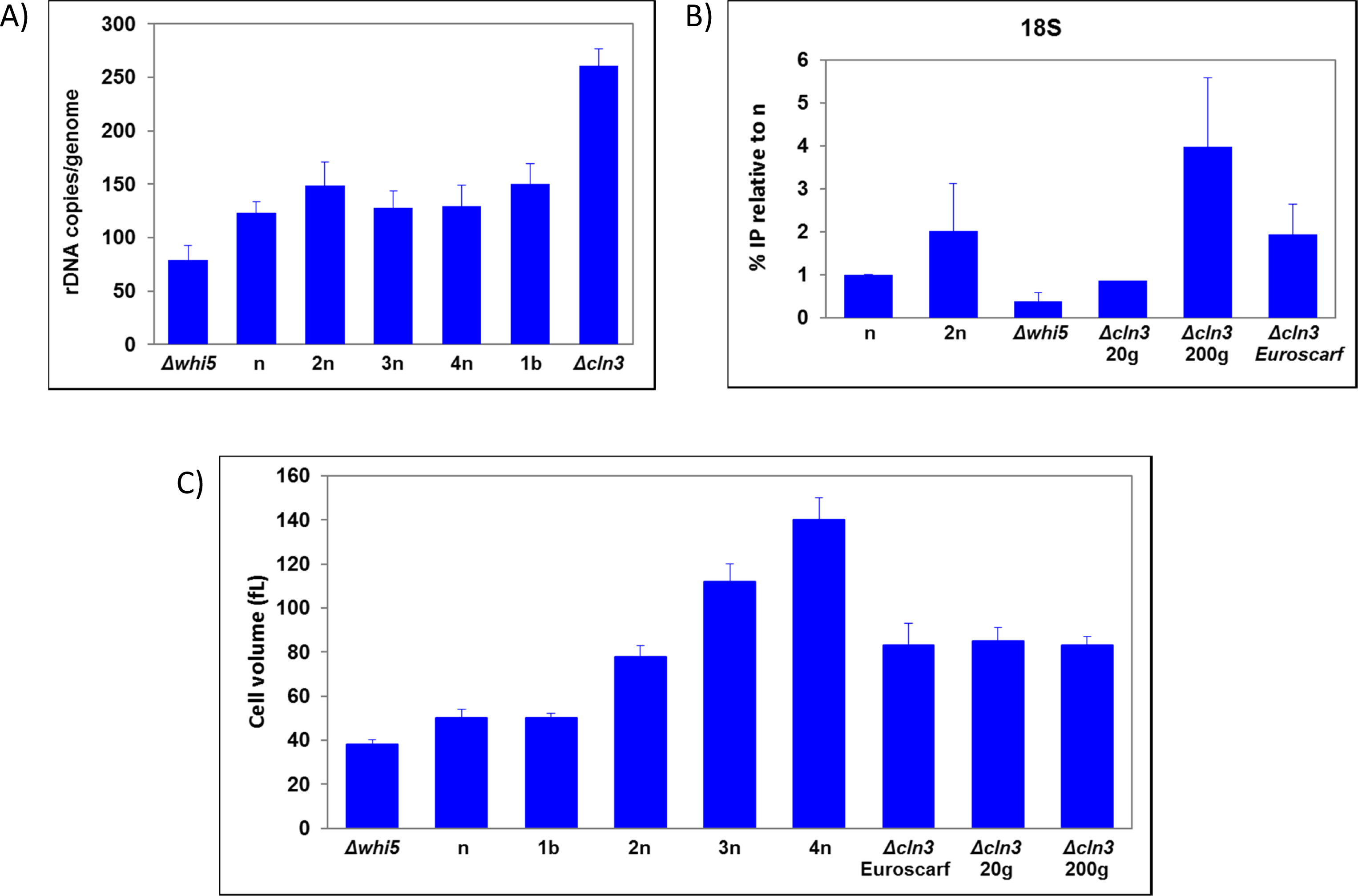
The number of rDNA repeats evolves in a *cln3* mutant to match the optimum value of the RNA pol I nTR required for its cell volume. The number of rDNA repeats varies in different strains (A) in a way that depends on ploidy, except for cell size haploid mutants (C). The RNA pol I nTR of the *cln3* mutants varies according to the rDNA repeat number, as determined by chromatin immunoprecipitation of the second largest RNA pol I subunit (A135) onto 18S (B). This figure is complementary to Figure 3.

**Supplementary Table 1.** Yeast strains used in this study.

| Strain | Genotype | Source | Ploidy | Doubling time (h) | Volume (fL) | Relative Volume |
| --- | --- | --- | --- | --- | --- | --- |
| **BY4741 (n)** | *MAT a, leu2Δ0, his3Δ1, met15Δ0, ura3Δ0* | Euroscarf | n | 1.7 | 50 ± 4 | 1 |
| **PY4993 (2n)** | *MAT a/α; ura3Δ0/ura3Δ0, leu2Δ0/leu2Δ0, his3Δ1/his3Δ1, met15Δ0/MET15, LYS2/lys2Δ0* | Pellman Lab | 2n | 1.7 | 78 ± 5 | 1.57 |
| **PY4997 (3n)** | *MAT a/a/α; ura3Δ0/ura3Δ0/ura3Δ0, leu2Δ0/leu2Δ0/leu2Δ0, his3Δ1/his3Δ1/his3Δ1, met15Δ0/MET15/MET15/, lys2Δ0/lys2Δ0/LYS2* | Pellman Lab | 3n | 1.7 | 112 ± 8 | 2.26 |
| **PY4996 (4n)** | *MAT a/a/α/α;;ura3Δ0/ura3Δ0/ura3Δ0/ura3Δ0, leu2Δ0/leu2Δ0/leu2Δ0/leu2Δ0, his3Δ1/his3Δ1/his3Δ1/his3Δ1, met15Δ0/met15Δ0/MET15/MET15/, lys2Δ0/lys2Δ0/LYS2/LYS2* | Pellman Lab | 4n | 1.7 | 140 ± 10 | 2.83 |
| **JCY0704** | BY4741 *whi5::LEU2* | JC Igual Lab | n | 1.9 | 38 ± 3 | 0.76 |
| **BQS2006 (Euroscarf Δcln3)** | Haploid strain derived from Y30366 diploid (Euroscarf). *MAT?*; *ura3Δ, leu2Δ, HIS3, MET15, lys2Δ, cln3Δ::KanMX4,* | our lab | n | 1.9 | 83 ± 10 | 1.68 |
| **BF305-15d** | *MATa, cln3::URA3-GAL1-CLN3, cln1::HIS3, cln2::TRP1, leu2, his3, ura3, trp1, ade1, arg5,6, met14* | M. Aldea Lab | n | nd | 60 ± 4 | 1.2 |
| **NOY408-1b** | *MAT a ade2-1 ura3-1 his3-11 trp1-1 leu2-3, 112 can1-100 pNOY102* | Kobayasi Lab | n | 2 | 43.7 ± 9 | 1.02* |
| **NOY408-1a** | *MAT a ade2-1 ura3-1 his3-11 trp1-1 leu2-3, 112 can1-100 rpa135:: LEU2 pNOY102* | Kobayasi Lab | n | nd | 59.54 ± 0 | 1.38* |
| **NOY408-1af** | *MAT a ade2-1 ura3-1 his3-11 trp1-1 leu2-3, 112 can1-100 rpa135:: LEU2 fob1::HIS3 pNOY102* | Kobayasi Lab | n | 3.2 | 66.89 ± 3 | 1.56* |

\*Relative volume values were obtained by considering BY4741 volume when growing in YPGal. Rest of volumes obtained from growth in YPD exponential phase. n.d. not determined

**Supplementary Table 2.**
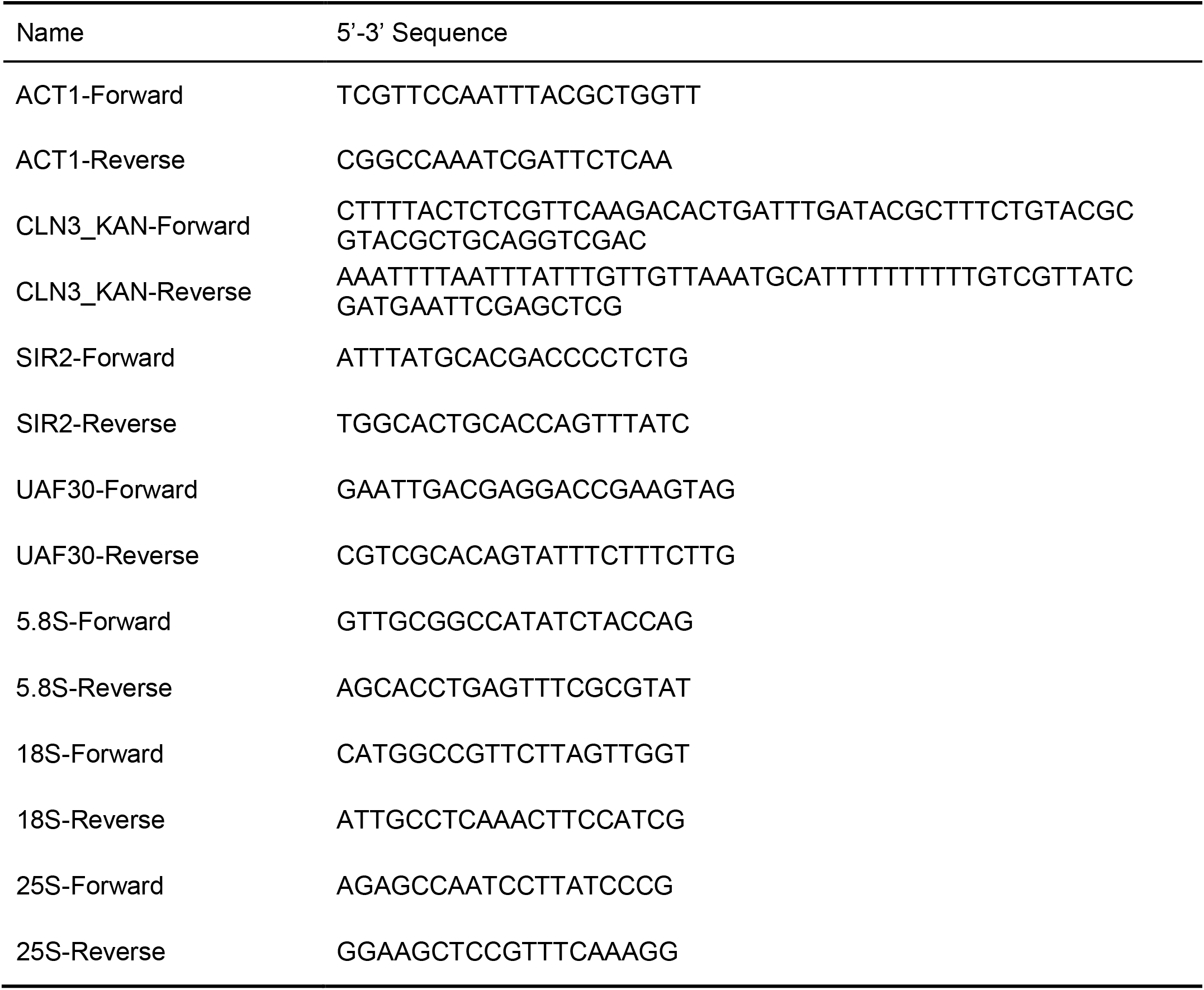
Oligonucleotides used in this study.

## Appendix: Integral Feedback

Consider the following system of chemical reactions which correspond to UAF molecules binding/unbinding to rDNA

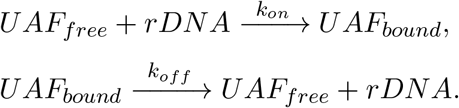

We assume that the total number of UAF molecules inside the cell scales with volume, and is given by *cV*, where *V* is the cell volume, and *c* is the UAF concentration. Then the number of bound UAF molecules bound to rDNA is given by

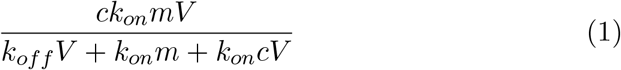

where *m* is the number of rDNA repeats. In the limit of strong binding of UAF to rDNA (*k*_*off*_ ≪ *k*_*on*_*c*), the concentration of free UAF molecules

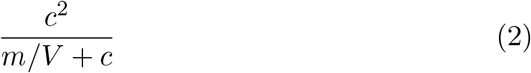

increases with increasing *V*. We assume that the production of SIR2 is a monotonically decreasing function of 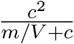 and given by 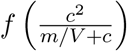. Let *x*(*t*) denote the level of SIR2, then it evolves as per the differential equation

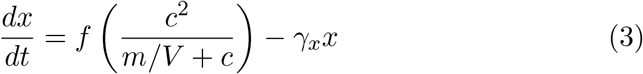

where *γ*_*x*_ is the SIR2 degradation rate. The number of rDNA repeats *m*(*t*) follows the differential equation

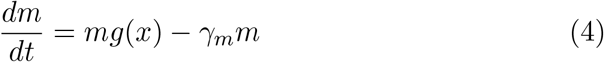

where *γ*_*m*_ and *g*(*x*) are rates of deletion and expansion of rDNA repeats, respectively. A key feature here is that *g*(*x*) is a monotonically decreasing function of SIR2 levels. The mathematical model given by (3)-(4) has been referred to in literature as *integral feedback control* (Aoki et al, Nature 2019) and it ensures that the the concentration of rDNA repeats is robustly maintained at a set point. To see this, consider equation (4) at steady-state, which gives the equilibrium SIR2 levels as

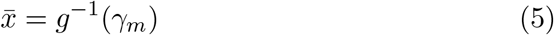

where *g*^−1^ denotes the inverse transformation of *g*. Now using (3) we can see that the steady-state rDNA repeat concentration 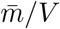 should satisfy

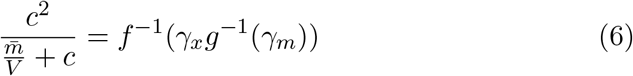

implying the setpoint

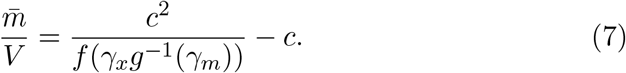

In essence, an increase in volume results in an increase in the concentration of free UAF molecule as per (2). This causes the SIR2 production rate, and its levels *x* to decrease, which in turn increases the rate of expansion

